# *ebony* affects pigmentation divergence and cuticular hydrocarbons in *Drosophila americana* and *D. novamexicana*

**DOI:** 10.1101/2020.03.05.977009

**Authors:** Abigail M. Lamb, Zinan Wang, Patricia Simmer, Henry Chung, Patricia J. Wittkopp

## Abstract

*Drosophila* pigmentation has been a fruitful model system for understanding the genetic and developmental mechanisms underlying phenotypic evolution. For example, prior work has shown that divergence of the *tan* gene contributes to pigmentation differences between two members of the virilis group: *Drosophila novamexicana*, which has a light yellow body color, and *D. americana*, which has a dark brown body color. Quantitative trait locus (QTL) mapping and expression analysis has suggested that divergence of the *ebony* gene might also contribute to pigmentation differences between these two species. Here, we directly test this hypothesis by using CRISPR/Cas9 genome editing to generate *ebony* null mutants in *D. americana* and *D. novamexicana* and then using reciprocal hemizygosity testing to compare the effects of each species’ *ebony* allele on pigmentation. We find that divergence of *ebony* does indeed contribute to the pigmentation divergence between species, with effects on both the overall body color as well as a difference in pigmentation along the dorsal abdominal midline. Motivated by recent work in *D. melanogaster*, we also used the *ebony* null mutants to test for effects of *ebony* on cuticular hydrocarbon (CHC) profiles. We found that *ebony* affects CHC abundance in both species, but does not contribute to qualitative differences in the CHC profiles between these two species. Additional transgenic resources for working with *D. americana* and *D. novamexicana*, such as *white* mutants of both species and *yellow* mutants in *D. novamexicana*, were generated in the course of this work and are also described. Taken together, this study advances our understanding of loci contributing to phenotypic divergence and illustrates how the latest genome editing tools can be used for functional testing in non-model species.

## 2 Introduction

Insect pigmentation is a well-studied trait that displays a variety of phenotypic differences within and between species (Kronforst et al. 2012; Wittkopp et al. 2003). These differences have evolved over a wide range of divergence times and in a great diversity of ecological contexts. Differences in insect pigmentation often appear to be ecologically relevant, correlating with geographic and climatic factors and playing a role in phenomena such as mate recognition, camouflage, thermoregulation, and water balance (True 2003; Wittkopp and Beldade 2009). Studies of pigmentation differences within the genus *Drosophila* have emerged as a productive model for studying the evolution of development, exploiting the diversity of phenotypes as well as genetic tools available for working with Drosophila and a long history of research into the genetic and biochemical mechanisms controlling pigmentation development (Wittkopp et al. 2003; Massey and Wittkopp 2016; Rebeiz and Williams 2017). Indeed, since the early 2000s, the genetic bases of dozens of pigmentation differences have been identified in varying levels of detail. Strikingly, in every case where a causal role has been directly attributed to a specific gene, the mechanism of change has been found to be a *cis*-regulatory change that affects gene expression rather than a change in the protein’s function (Massey and Wittkopp 2016). These case studies have also identified multiple independent instances of divergent expression for some pigmentation genes, suggesting that these genes are particularly tractable routes for the evolution of pigmentation in this genus (Massey and Wittkopp 2016).

Changes in *cis*-regulatory sequences are thought to be a common mechanism of developmental evolution because they tend to be less pleiotropic than changes in protein function (Wray et al. 2003; Carroll 2005). For example, a *cis*-regulatory change might alter a gene’s expression in only a single tissue or a single point in development whereas changing its protein function is expected to impact the organism everywhere that protein is expressed. Genes controlling pigmentation development in Drosophila might be especially likely to evolve using this mechanism because the proteins encoded by these genes are also required for other biological functions. For example, genes required for pigment synthesis have also been shown to affect mating success, circadian rhythm, vision, and innate immunity (Massey et al. 2019a; Suh and Jackson 2007; True et al. 2005; Nappi and Christensen 2005; Takahashi 2013; Wittkopp and Beldade 2009). The pigmentation biosynthesis genes *ebony* and *tan* have also been found to affect the profiles of cuticular hydrocarbons on adult flies, which are hydrophobic lipids on the surface of insect cuticle that are involved in chemical communication, mate recognition, and water balance (Massey et al. 2019b; Chung and Carroll 2015; Chung et al. 2014).

Here, we investigate genetic changes contributing to the evolution of novel body color in *D. novamexicana*. This species has evolved a much lighter and more yellow body color than its sister species *D. americana* during the approximately 400,000 years since these species diverged from their most recent common ancestor (Figure 1, Caletka and McAllister 2004; Morales-Hojas et al. 2008). *D. novamexicana* and *D. americana* show signs of reproductive isolation (Ahmed-Braimah and McAllister 2012; Patterson and Stone 1949), but they are interfertile and can produce viable, fertile F_1_ hybrids in the laboratory, allowing genetic analysis (Wittkopp et al. 2003; Wittkopp et al. 2009). Prior genetic mapping has identified two quantitative trait loci (QTL) that together account for ~87% of the pigmentation difference between *D. novamexicana* and *D. americana* (Wittkopp et al. 2009). Fine mapping and transgenic analysis revealed that the QTL of smaller effect was driven by divergence at *tan* (Wittkopp et al. 2009), a gene that encodes a hydrolase that catalyzes the conversion of N-B-alanyl dopamine (NBAD) to dopamine, a precursor for dark melanin pigment (True et al. 2005). The QTL of larger effect was linked to an inverted region containing the candidate gene *ebony*, but the presence of the inversion prevented fine mapping to separate the effects of *ebony* from linked loci (Wittkopp et al. 2009). *ebony* encodes a synthetase that catalyzes the conversion of dopamine into NBAD, a precursor for light yellow pigments (Koch et al. 2000), which is the opposite of the reaction catalyzed by Tan. *ebony* has also been shown to have expression differences between *D. novamexicana* and *D. americana* caused by *cis-*regulatory divergence (Cooley et al. 2012).

**Figure 1.**
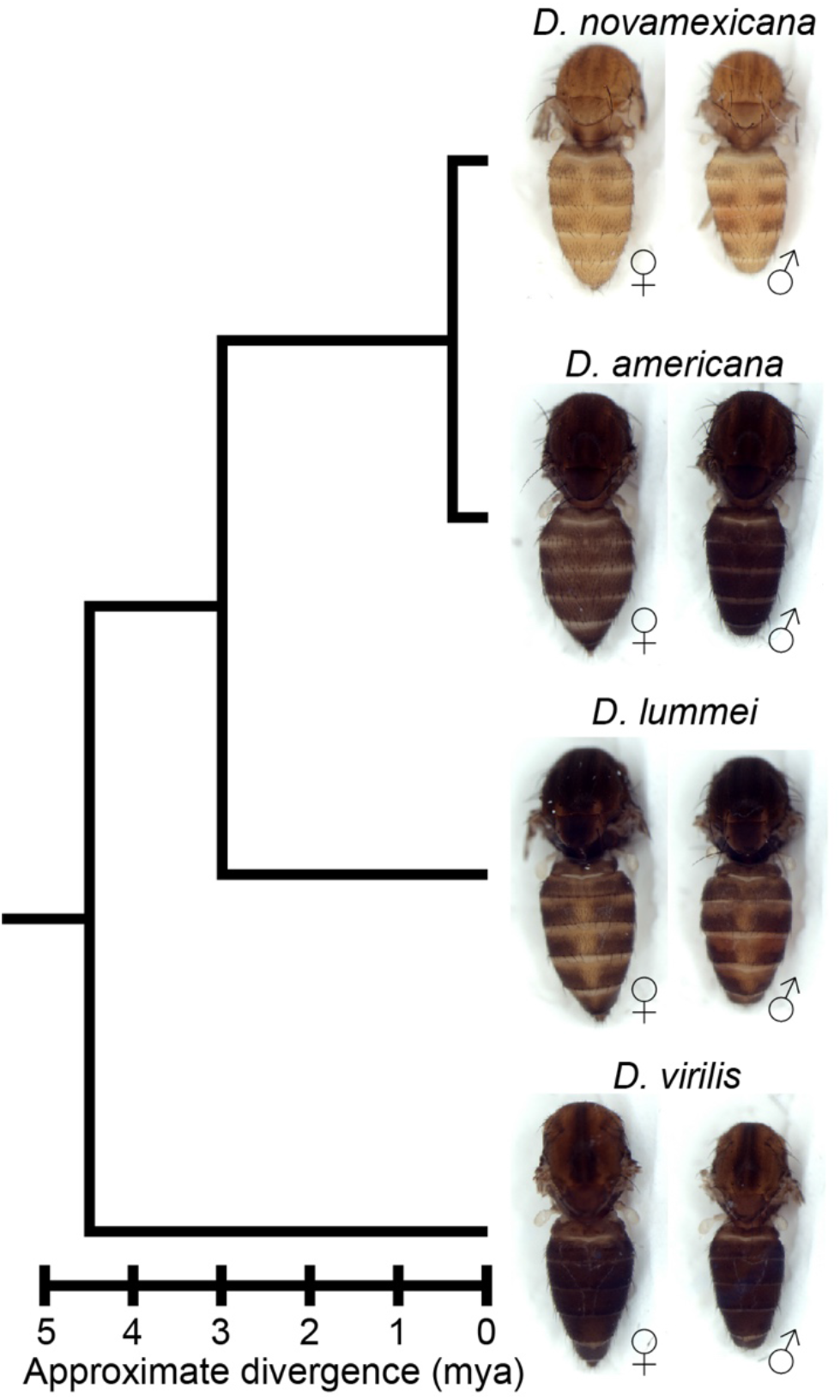
*D. novamexicana* shows divergent body color within the virilis group. Phylogenetic relationships with estimated divergence times (Caletka and McAllister 2004; Cooley et al 2012) are shown for *D. novamexicana*, *D. americana*, *D. lummei*, and *D. virilis*. For each species, a dorsal view of the thorax and abdomen is shown for females (left) and males (right), with heads, wings, and legs removed.

Despite these data suggesting that *ebony* contributes to pigmentation divergence between *D. novamexicana* and *D. americana*, the phenotypic effects of sequence divergence at *ebony* have not been demonstrated. Here, we show that divergence at *ebony* does indeed contribute to pigmentation divergence between these two species. We use CRISPR/Cas9 genome editing to mutate *ebony* in both species and use these mutant genotypes to directly test *ebony*’s contribution to pigmentation divergence through reciprocal hemizygosity testing (Stern 2014). We find that the *D. novamexicana ebony* allele causes lighter pigmentation throughout the body than the *D. americana ebony* allele. We also find that allelic divergence at *ebony* is primarily responsible for a spatial difference in abdominal pigmentation between these species: the *D. novamexicana ebony* allele causes the absence of dark melanin along the dorsal midline of the abdomen seen in *D. novamexicana*. Finally, we show that *ebony* affects the cuticular hydrocarbon (CHC) profiles in *D. americana* and *D. novamexicana*, but does not contribute to the qualitative differences in CHC profiles seen between species. Taken together, our data show the power of using CRISPR/Cas9 genome editing to test functional hypotheses about evolutionary mechanisms. In addition, resources generated and lessons learned in the course of this work are expected to help other researchers perform CRISPR/Cas9 genome editing in *D. americana, D. novamexicana* and other Drosophila species.

## 3 Materials and Methods

### 3.1 Fly stocks and husbandry

The following fly lines were used in this study: *D. americana* “A00” (National Drosophila Species Stock Center number 15010-0951.00), *D. novamexicana* “N14” (National Drosophila Species Stock Center number 15010-1031.14), *D. lummei* (National Drosophila Species Stock Center number 15010-1011.08), *D. virilis* (National Drosophila Species Stock Center number 15010-1051.87), *D. melanogastery^1^ M{w[+mC]=nos-Cas9.P}ZH-2A w*\* (Bloomington Drosophila Stock Center number 54591), and *D. melanogaster* Canton-S. All flies were reared on standard cornmeal medium at 23-25°C with a 12:12 hour light:dark cycle.

### 3.2 Transgenesis and CRISPR mutant generation in *D. americana* and *D. novamexicana*

To the best of our knowledge, prior to this work, the only transformation of *D. americana* or *D. novamexicana* resulted from the insertion of a piggyBac transgene (Wittkopp et al. 2009). We therefore first used the CRISPR/Cas9 system to generate *white* mutants in both species to test the feasibility of CRISPR genome modification and to create lines that are easier to screen for common transformation markers that drive expression of fluorescent proteins or restore red pigmentation in the eyes by restoring *white* function. We successfully generated *white* mutant N14 and A00 lines used as transgenic hosts for future work, by injecting single guide RNAs (sgRNAs) targeting coding sequences in *white* conserved between *D. novamexicana* and *D. americana* in the second and third exons and screening for the loss of red eye pigment in male offspring of injected females (Supplementary Figure 1); *white* is on the X chromosome and thus only present in a single copy in males. These same guide RNAs were also used in *Drosophila virilis* to cut *white* and integrate an attP landing site potentially useful for site-directed transgene insertion (J. Lachoweic and P.J. Wittkopp, unpublished data), although the PhiC31 system does not seem to work well in *D. virilis* (Stern et al. 2017). For all CRISPR experiments, sgRNAs were in-vitro transcribed from DNA templates using Invitrogen T7 MEGAscript Transcription Kit according to protocol described by Bassett et al. 2013. Oligonucleotides used to generate sgRNAs are listed in Supplementary Table 1. After transcription, sgRNAs were purified using RNA Clean and Concentrator 5 kit (Zymo Research), eluted with nuclease-free water, and quantified with Qubit RNA BR Assay Kit (Thermo Fisher Scientific). For CRISPR injections, sgRNAs were mixed with purified Cas9 protein (PNA Bio #CP01) with a final injected concentration of 0.05% phenol red to visualize the injection mix. CRISPR injections were performed in-house, using previously described methods (Miller et al. 2002).

To try to increase efficiency of CRISPR mutagenesis in these species, we next sought to generate transgenic lines expressing Cas9 in the germlines of *white* mutant *D. americana* (A00) and *D. novamexicana* (N14) flies using piggyBac transgenesis (Horn and Wimmer 2000). Based on prior reports that the *nanos (nos)* promoter and 3’UTR drive expression in the germline of *Drosophila virilis* (Holtzman et al. 2010), a close relative of *D. americana* and *D. novamexicana*, we amplified the *nos-Cas9-nos* transgene from the pnos-Cas9-nos plasmid (Addgene #62208; Port et al. 2014) using Phusion High Fidelity Polymerase (NEB) with tailed primers and cloned the amplicon into pBac{3XP3-ECFPafm} (Horn and Wimmer 2000) digested with AscI and Bsu36I restriction enzymes using Gibson Assembly Master Mix (NEB). Primers are included in Supplemental Table 1. We confirmed the insert was correctly incorporated and free of PCR-induced errors by Sanger sequencing. We sent the *white* mutant lines of *D. americana* A00 and *D. novamexicana* N14 that we generated to Rainbow transgenic services for piggyBac transgenesis (www.rainbowgene.com) and screened offspring of injected adults for expression of the enhanced cyan fluorescent protein (ECFP) in the eye using a Leica MZ6 stereoscope equipped with a Kramer Scientific Quad Fluorescence Illuminator. Transformants were obtained from injections into *D. novamexicana* (N14) (PCR verified), but not from injections into *D. americana* (A00), despite multiple attempts.

All subsequent CRISPR injections in *D. novamexicana* were performed using flies homozygous for the *nos-Cas9-nos* transgene, some with and some without the inclusion of commercially available Cas9 protein in the injection mix. CRISPR mutants were only obtained from injections containing the commercially available Cas9 protein, however, suggesting that the *nos-Cas9-nos* transgene might not drive expression of Cas9 in the germline of *D. novamexicana*. To test this hypothesis, we used western blotting to examine Cas9 protein expression in 3 transformed *D. novamexicana* N14 lines with independent insertions of the piggyBac transgene and in a *D. melanogaster* transgenic line carrying the original *pnos-Cas9-nos* transgene (Bloomington Drosophila Stock Center line 54591, transformed with Addgene plasmid #62208, Port et al. 2014). These experiments showed that the *nos-Cas9-nos* transgene in *D. novamexicana* N14 flies does not express Cas9 protein in the ovaries (Supplementary Figure 2). This conclusion was further supported when injection of sgRNAs targeting the *yellow* gene into the *D. novamexicana* line carrying the *nos-Cas9-nos* transgene also only produced *yellow* mutants when the Cas9 protein was co-injected with the sgRNAs (Supplementary Figure 3). Ability of the *nos* promoter to drive germine expression in the closely related species *D. virilis* has also been found to be variable among transgenic lines (Hannah McConnell, Aida de la Cruz, and Harmit Malik, personal communication), suggesting that other promoters should be used in the future to drive reliable germline expression in the virilis group.

To generate *ebony* mutant *D. americana* (A00) and *D. novamexicana* (N14), we synthesized five sgRNAs targeting conserved sites in the first coding exon of *ebony*. Because *ebony* is located on an autosome and *ebony* loss-of-function mutant alleles are generally considered recessive in *D. melanogaster* (Thurmond et al. 2019), we did not expect to be able to identify *ebony* mutants by simply screening progeny of injected flies for mutant phenotypes as we did for *white* and *yellow*. We therefore co-injected a donor plasmid containing the sequence of an eye-specific red fluorescent protein marker (3XP3-RFP) flanked by *ebony* sequences that could be inserted into *ebony* via homology-directed repair and used to screen for *ebony* mutants. Although we observed RFP expression in larvae injected with the homology-directed repair donor fragment, indicating that the reporter gene was functional in these species, injected individuals did not produce any offspring with red fluorescent eyes, suggesting that the donor plasmid was not integrated in the germline of injected individuals. Because non-homologous end joining occurs more frequently than homology directed repair following double-strand breaks (Liu et al. 2018), we also tried to identify flies that might be heterozygous for an *ebony* mutant allele by closely inspecting all offspring of injected (G0)flies for any subtle changes in pigmentation. Specifically, we collected and mated (G1) offspring of injected flies with any noticeably darker pigmentation, keeping them grouped by G0 parent of origin. As further described in the results, we were ultimately able to identify homozygous *ebony* mutants among progeny from these G_1_ x G_1_ crosses of relatively dark flies derived from two independent *D. novamexicana* G_0_ flies and one *D. americana* G_0_ fly. Sanger sequencing these flies confirmed they were homozygous for *ebony* alleles containing deletions. We then crossed the mutated *ebony* alleles back into wild-type backgrounds of each parental species to generate homozygous *ebony* mutant lines with wild-type red eyes.

### 3.3 Western blotting

For *ebony* western blotting, proteins were extracted from stage P14/15 pupae, identified by the following characteristics: black pigmentation present in wings and bristles, meconium visible in abdomen (Cooley et al. 2012). For each sample, five pupae were homogenized in 100uL of homogenization buffer (125 mM Tris pH 6.8, 6% SDS, 2.5X Roche cOmplete protease inhibitor cocktail, EDTA-free), then centrifuged for 15 minutes at 15000rcf, and the supernatant transferred to a fresh tube with an equal volume of 2x Laemmli buffer (125 mM Tris pH 6.8, 6% SDS, 0.2% glycerol, 0.25% bromophenol blue, 5% Beta-mercaptoethanol).

For Cas9 western blotting, protein was extracted from ovaries dissected in ice cold PBS from the following lines: untransformed N14 *white* mutants (host line), three independently transformed lines of N14 *white* carrying the pBac{3XP3-ECFPafm-nosCas9nos} transgene, transgenic *D. melanogaster* carrying the *pnos-Cas9-nos* transgene, and wild-type (Canton-S) *D. melanogaster*. For *D. novamexicana* samples, we collected ovaries from 10 sexually mature flies, whereas for *D. melanogaster* samples, we collected ovaries from 18 sexually mature flies. Different numbers of flies were used for the two species because of differences in body size. In each case, ovaries were placed into microcentrifuge tubes on ice, spun down briefly in a tabletop centrifuge, and excess PBS was removed and replaced with 20uL of homogenization buffer. Samples were then treated as described for *ebony* western blots above. A positive control Cas9 sample was made by diluting purified Cas9 protein (PNA Bio CP01) in homogenization buffer, and mixing with 2X Laemmli buffer to a final concentration of 2.5ng/uL.

Samples were heated at 95°C for 10 minutes before loading into 7.5% Mini-PROTEAN® TGX™ Precast Protein Gels (BioRad) and running at 150V for approximately 90 minutes at 4°C in 1X tris-gylcine running buffer. Separate gels were run for *ebony* and Cas9 blots. Samples were loaded in the following volumes: 35uL per pupa sample, 30uL per ovary sample, 10uL of Cas9 positive control (25ng protein), 5uL PageRuler prestained protein ladder (Thermo Scientific). Gels were transferred onto PVDF membrane in tris-glycine transfer buffer, 10% MeOH, 0.01% SDS at 100V for 1 hour with stirring on ice at 4°C. Membranes were blocked in 3% nonfat dry milk in TBST for 30 minutes at RT with shaking, then divided in half using the prestained ladder as a guide just below the 100kDa mark for the Cas9 membrane and just below the 70kDa mark for the *ebony* membrane. The lower molecular weight halves of the membranes were placed in solutions containing primary antibodies to detect the protein used as a loading control (tubulin or lamin), whereas the halves of the membranes containing the higher molecular weight proteins were placed in solutions containing primary antibody solutions against the protein of interest (Ebony or Cas9), each diluted in 3% nonfat dry milk in TBST. In all cases, membranes were incubated with the primary antibodies overnight at 4°C. Primary antibody solutions for *ebony* included rabbit anti-*ebony* 1:300 (Wittkopp et al. 2002) and rabbit anti-alpha tubulin 1:5000 (Abcam ab52866) as a loading control. Primary antibody solutions for Cas9 included mouse anti-Cas9 1:1000 (Novus NBP2-36440) and mouse anti-lamin 1:200 (DHSB adl67.10) as a loading control. Membranes were washed in TBST and transferred to secondary antibody solutions diluted in 3% nonfat milk in TBST for 2 hours at RT. The following secondary antibodies were used: donkey anti-rabbit HRP 1:5000 (Amersham na934) or goat antimouse HRP 1:5000 (abcam ab97023). Membranes were finally washed in TBST and developed with SuperSignal West Pico Chemiluminescent Substrate (Thermo Scientific) and imaged using a Licor Odyssey FC imaging system.

### 3.4 Fly crosses for reciprocal hemizygosity testing and cuticular hydrocarbon analysis

To generate F_1_ hybrids carrying only one (*D. americana* or *D. novamexicana)* functional *ebony* allele, wild-type and *ebony* mutant flies from each species were collected as virgins and aged in vials for at least 12 days to reach sexual maturity and verify virgin female status by absence of larvae. Crosses were all set on the same batch of food on the same day and placed at 25°C. For most crosses, 4 virgin females and 4 males were used; however, 8 virgin females and 8 males were used in interspecific crosses with *D. novamexicana* females because of reduced mating success in these crosses. After 3 days, adult flies from these crosses that would be used for cuticular hydrocarbon (CHC) analysis were transferred to new vials with a fresh batch of food. Offspring from the first set of vials were used for imaging and pigmentation analysis, while offspring from the second set of vials were used for CHC analysis. Flies used for pigmentation phenotyping were aged 5-7 days after eclosion and preserved in 10% glycerol in ethanol before imaging (Wittkopp et al. 2011).

### 3.5 Imaging of fly phenotypes

Insect specimens were imaged using a Leica DC480 camera attached to a Leica MZ16F stereoscope equipped with a ring light attachment and Leica KL 1500 LCD lamp. Images were captured using Leica DC Twain software version 5.1.1 run through Adobe Photoshop CS6 version 13.0 X32. Prior to imaging, pupal cases and wings were mounted on slides in PVA mounting medium (BioQuip). Thorax, abdomen, and whole-body specimens were prepared from age-matched, preserved flies as described in the previous section. For imaging, thorax, abdomen, and whole-body specimens were submerged in 100% ethanol in custom wells composed of white oven-cured polymer clay (Sculpey).

Because the color of specimens spanned a wide range across genotypes, exposure was optimized for each sample type (e.g. whole body, thorax, abdomen, wing, pupal case) individually by placing specimens from the two phenotypic extremes in the same frame and adjusting exposure to avoid over-exposing the lightest flies while capturing as much detail as possible from the darkest flies. Exposure time, lighting, white balance, background, and zoom were kept identical across all images of single tissue type. Minor color adjustments to improve visibility of phenotypes were performed simultaneously across all raw images of the same sample type in a single combined document using Photoshop CC 2019, ensuring that all images presented for direct comparisons were adjusted identically.

### 3.6 Cuticular hydrocarbon analyses

CHCs for each cross were extracted from five 5-day-old females by soaking the flies for 10 mins in 200 μl hexane containing hexacosane (C26; 25 ng/ul) as an internal standard. Eight replicates were prepared for each cross. Extracts were directly analyzed by the GC/MS (7890A, Agilent Technologies Inc., Santa Clara, CA) coupled with a DB-17ht column 30 m by 0.25 mm (i.d.) with a 0.15 μm film thickness (Agilent Technologies Inc., Santa Clara, CA). Mass spectra were acquired in Electron Ionization (EI) mode (70 eV) with Total Ion Mode (TIM) using the GC/MS (5975C, Agilent Technologies Inc., Santa Clara, CA). The peak areas were recorded by MassHunter software (Agilent Technologies Inc., Santa Clara, CA). Helium was the carrier gas at 0.7 ml/min and the GC thermal program was set as follows: 100 °C for 4 min, 3 °C/min to 325 °C. Straight-chain compounds were identified by comparing retention times and mass spectra with authentic standard mixture (C6-C40) (Supelco® 49452-U, Sigma-Aldrich, St. Louis, MO). Methyl-branched alkanes, alkenes, dienes and trienes were then identified by a combination of their specific fragment ions on the side of functional groups (methyl branch or double bonds) and retention times relative to linear-chain hydrocarbon standards. Each individual CHC peak was quantified by normalizing its peak area to the peak area of the internal C26 standard, converting each CHC peak area to ng/fly using the known internal standard concentration of 1000 ng/fly. Welch’s *t*-tests with a Benjamini-Hochberg correction for multiple testing (Benjamini and Hochberg 1995) were used to compare CHC amounts between pairs of genotypes. To compare the effects of *ebony* loss of function on different chain-lengths of CHCs, eight biological replicates of homozygous *ebony* null measurements were divided by the mean measurement of the eight replicates of the matched *ebony* heterozygote for each individual CHC. The ratio of *ebony* null to heterozygote CHC abundance was plotted against CHC chain length. The relative effects of *D. americana* versus *D. novamexicana ebony* in a common F_1_ hybrid background (described as F_1_[*e^A^/e^-^*] and F_1_[*e^N^/e*^-^], respectively) were also compared in this manner, with the replicates of the F_1_[*e^A^*/*e*^-^] divided by the mean F_1_[*e^N^/e*^-^] measurement for each CHC. We used Spearman’s rank correlation (Spearman’s rho) to test the relationship between CHC chain length and the effect of *ebony* on CHC abundance. The threshold for statistical significance was set at *alpha* = 0.05 for all tests. Datafile and R code used for this analysis are provided in Supplementary Files 1 and 2, respectively

## 4 Results and Discussion

The reciprocal hemizygosity test is a powerful strategy for identifying genes with functional differences that contribute to phenotypic divergence (reviewed in Stern 2014). This test is performed by comparing the phenotypes of two hybrid genotypes that are genetically identical except for which allele of the candidate gene is mutated. Any phenotypic differences observed between these two genotypes are attributed to divergence of the candidate gene. Applying this test to identify functional differences between species requires loss-of-function (null) mutant alleles in both species and the ability for the species to cross and produce F_1_ hybrids. Consequently, in order to use this strategy to test *ebony* for functional divergence between *D. novamexicana* and *D. americana*, we first needed to generate *ebony* null mutant alleles in both species.

### 4.1 Generating *ebony* mutants in *D. americana* and *D. novamexicana* using CRISPR/Cas9

We generated *ebony* null mutants in *D. novamexicana* and *D. americana* by using CRISPR/Cas9 to target double-strand breaks to five conserved sites within the first coding exon of *ebony*. As described more fully in the Methods section, we injected embryos of *white* mutants from both species with purified Cas9 protein and sgRNAs targeting all five sites simultaneously. To make it easier to identify *ebony* mutant alleles, we also injected a donor plasmid that would allow homology directed repair to integrate a transgene expressing red fluorescent protein in the fly’s eyes, but no progeny of injected flies were observed to express this transformation marker. We reasoned that *ebony* mutants might still have been generated by non-homologous end-joining, however, and thus also searched for *ebony* mutants by looking for changes in body pigmentation.

In *D. melanogaster, ebony* loss-of-function mutants have a much darker appearance than wild-type flies because they are unable to produce yellow sclerotin, causing an increase in production of black and brown melanins (Wittkopp et al. 2002). *D. melanogaster ebony* mutant alleles are commonly described as recessive to wild-type *ebony* alleles (Thurmond et al. 2019); however, in some genetic backgrounds, flies heterozygous for an *ebony* mutant allele are slightly darker than wild-type flies (Thurmond et al. 2019). Because *D. novamexicana* has such a light yellow body color (Figure 2A), we thought it possible that flies heterozygous for an *ebony* mutant allele might also show a detectable darkening of pigmentation; we were less optimistic about being able to detect heterozygous *ebony* mutants based on pigmentation in *D. americana* because its wild-type pigmentation is already very dark (Figure 2C). Nonetheless, we sorted through the progeny of injected *D. novamexicana* and *D. americana* flies, isolating any individuals that seemed to have darker pigmentation than their siblings and allowing these relatively dark flies to freely mate in vials segregated by injected parents, keeping individual “founder” mutations separate.

**Figure 2.**
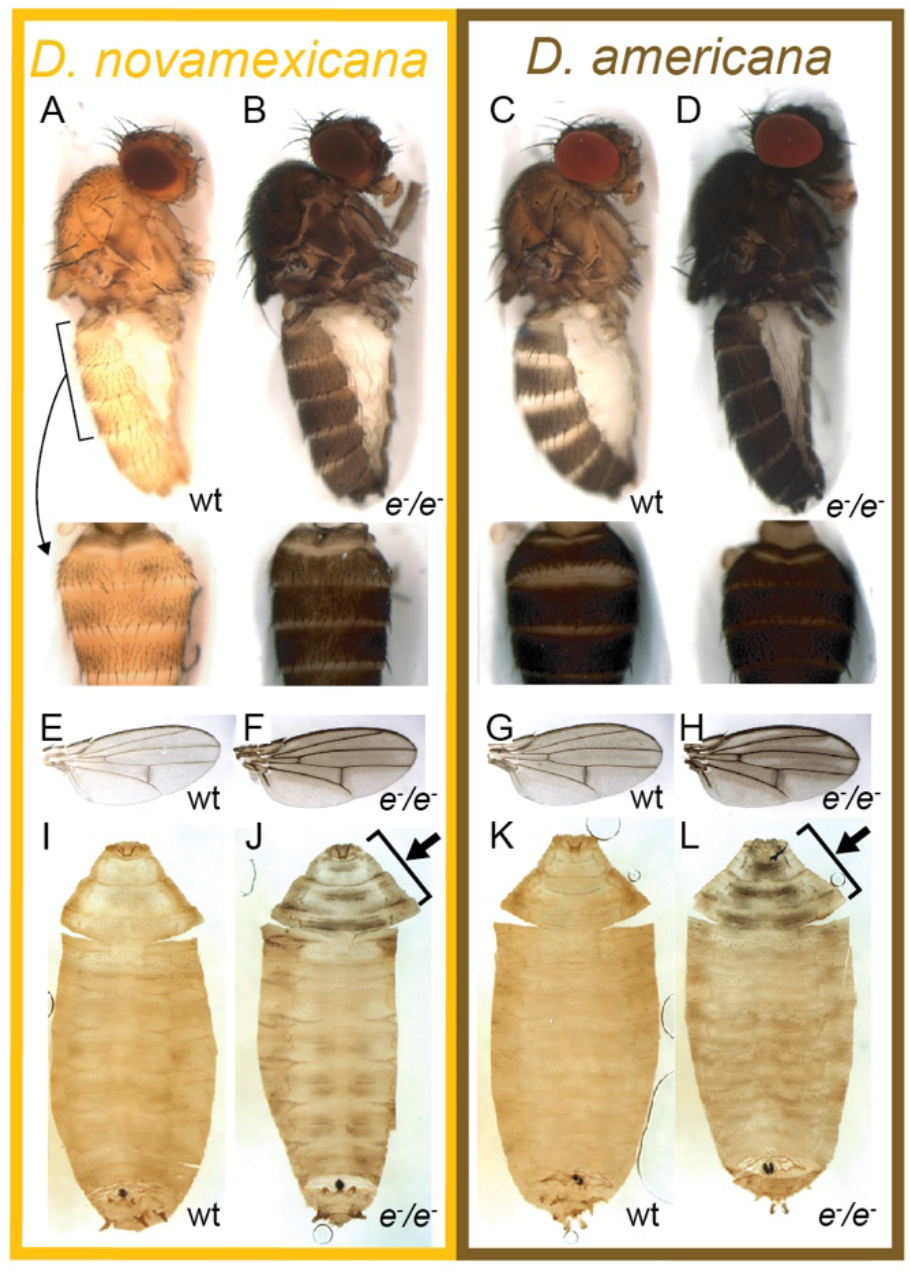
Ebony affects body, wing, and pupal pigmentation in *D. novamexicana* and *D. americana*. (**A-D**) Adult body pigmentation is shown from a lateral view (top) and dorsal abdominal view (segments A2-A4, bottom) for (**A**) *D. novamexicana*, (**B**) *D. novamexicana ebony* null mutants, (**C**) *D. americana*, and (**D**) *D. americana ebony* null mutants. (**E-H**) Adult wing pigmentation is shown for (**E**) *D. novamexicana*, (**F**) *D. novamexicana ebony* null mutants, (**G**) *D. americana*, and (**H**) *D. americana ebony* null mutants. (**I-L**) Pigmentation of pupal cases is shown for (**I**) *D. novamexicana*, (**J**) *D. novamexicana ebony* null mutants, (**K**) *D. americana*, and (**L**) *D. americana ebony* null mutants. Arrows in (**J**) and **(L)** highlight the most prominent areas with dark pigmentation in *ebony* mutants.

Two of the vials of darker pigmented *D. novamexicana* flies produced pupae with an unusual black pattern on the anterior end of the pupal case (Figure 2J). We moved these pupae to new vials and found that they developed into adults with the much darker than wild-type body color expected for homozygous *ebony* mutants in *D. novamexicana* (Figure 2 A,B). Because pigmentation of the pupal case is very similar between *D. novamexicana* and *D. americana* (Figure 2I,K, Ahmed-Braimah and Sweigart 2015), we also searched for pupae with similar pigmentation marks in the vials containing progeny of darker flies descended from injected *D. americana*. We found such pupae in one of the vials (Figure 2L). Flies emerging from these pupal cases also showed darker pigmentation than wild-type *D. americana* (Figure 2C,D), as expected for homozygous *ebony* mutants, but this difference was much more subtle than in *D. novamexicana* (Figure 2A,B). Flies from both species emerging from pupal cases with abnormal pigmentation also showed increased levels of dark melanins in wings in a pattern similar to that seen in *D. melanogaster ebony* mutants (Figure 2E-H, Wittkopp et al. 2002), further suggesting that they were homozygous for *ebony* mutant alleles. Crossing putative homozygous *ebony* mutants from the same species to each other resulted in true-breeding lines of *D. americana* and *D. novamexicana* presumed to be homozygous for *ebony* mutant alleles.

To determine whether these true-breeding lines were indeed homozygous for *ebony* mutant alleles, we used Sanger sequencing to search for changes in the *ebony* sequence in the region targeted for double strand breaks with CRISPR/Cas9. We found that the presumed *ebony* mutant lines of both species harbored deletions corresponding to the locations of sgRNA target sites in the first coding exon, with the two *D. novamexicana* mutant lines carrying deletions of 7 and 10 bases and the *D. americana* mutant line carrying a deletion of 46 bases (Figure 3A). Each of these mutations is expected to cause frameshifts, leading to multiple early stop codons. Further experiments described in this study using *D. novamexicana ebony* mutants were conducted with the 10 base deletion line, and any further description of *ebony* null *D. novamexicana* refers to this line.

**Figure 3.**
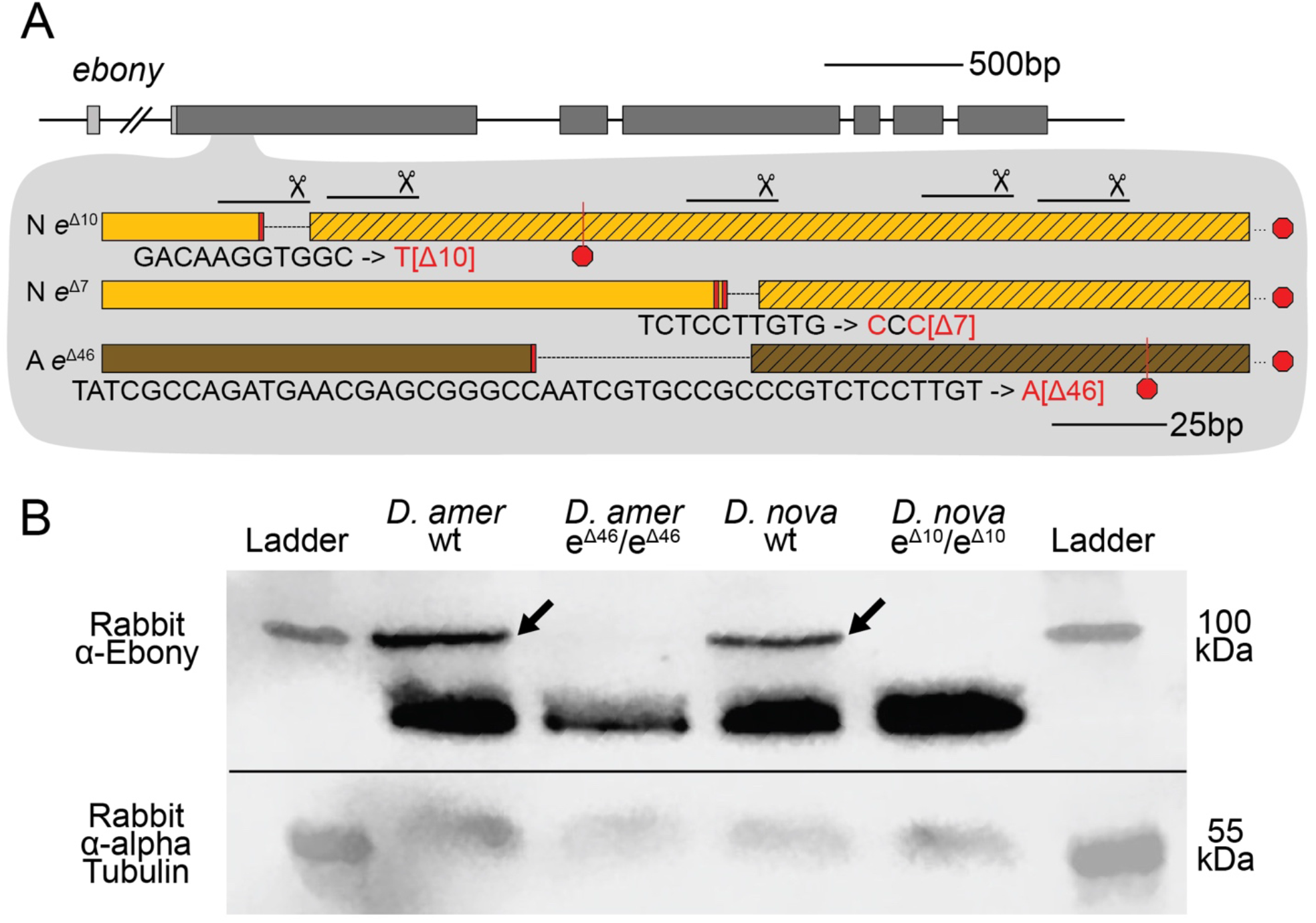
CRISPR/Cas9-induced mutations created null alleles of the *D. novamexicana* and *D. americana ebony* genes. (**A**) A schematic of the *ebony* gene is shown with grey boxes indicating exons; coding sequence is indicated in the darker shade of grey. Locations of the five guide RNAs targeting the second exon of *ebony* are shown with solid lines below scissor symbols. Mutations observed in the two *ebony* mutants (*e^Δ10^* and *e^Δ7^*) isolated in *D. novamexicana* (“N”) and the one *ebony* mutant (*e^Δ46^*) isolated in *D. americana* (“A”) are shown. All three alleles included deletions that caused frameshifts. (**B**) Western blotting showed that the *D. americana e^Δ46^* and *D. novamexicana e^Δ10^* mutants (lanes 2 and 4, respectively) lacked a ~100 kDa protein (arrows) recognized by an antibody raised against *D. melanogaster* Ebony protein (Wittkopp et al. 2002) that is present in wild-type (wt) *D. americana* and *D. novamexicana* (lanes 1 and 3, respectively). Relative abundance of total protein loaded into each lane can be seen by the relative intensities of the shorter proteins also detected by the Ebony antibody (Wittkopp et al. 2002) as well as the relative intensities of ~55kDa bands detected by an antibody recognizing alpha Tubulin (Abcam ab52866). The solid black line shows where the membrane was cut prior to incubation with primary antibodies during the western blotting procedure; the top half was incubated with anti-Ebony antibodies whereas the bottom half was incubated with anti-Tubulin antibodies. The two halves were realigned by hand for imaging, using the shape of the cut and the ladder staining as a guide. An un-annotated image of this blot is shown in Supplementary Figure 4.

To further assess whether these mutations caused null alleles, we used western blotting to examine the expression of the Ebony protein during late pupal stages when adult pigmentation is developing and the *ebony* gene is expressed in the developing abdomen (Wittkopp, et al. 2002; Cooley et al. 2012). We performed western blots on protein extracts from P14/P15 stage pupae of both wild-type and homozygous *ebony* mutant flies of both *D. americana* and *D. novamexicana* using an antibody against *D. melanogaster ebony* (Wittkopp et al. 2002). This antibody recognizes a 94 kDa protein consistent with the predicted molecular weight of Ebony in pupal protein extracts from wild-type lines of both *Drosophila melanogaster* and *Drosophila biarmipes*, but does not produce a 94kDa band in pupal protein extracts of either *e1* or *In(3R)eAFA ebony* mutant lines of *D. melanogaster* (Wittkopp et al. 2002). Wild-type extracts of both *D. americana* and *D. novamexicana* produced presumptive Ebony bands while extracts from flies homozygous for *ebony* deletions did not produce a 94 kDa band for either species (Figure 3B). The nature of the frameshift deletions as well as the western blot evidence together show that these *ebony* mutations cause null alleles.

### 4.2 *ebony* divergence contributes to body color differences between *D. novamexicana* and *D. americana*

We used the homozygous *ebony* mutant *D. novamexicana* and *D. americana* lines to perform a reciprocal hemizygosity test by crossing *ebony* mutant *D. novamexicana* (*e*^-^/*e*^-^) to wild-type *D. americana* (*e^A^/e^A^*) and *ebony* mutant *D. americana (*e*^-^/*e*^-^)* to wild-type *D. novamexicana* (*e^N^/e^N^*) (Figure 4A). In order to observe the effects of the two species’ *ebony* alleles in the presence of each species X chromosome, we conducted sets of reciprocal crosses (i.e., swapping the genotypes of the male and female parents). Female F_1_ hybrids from reciprocal crosses are genetically identical except for the parent of origin of their one functional *ebony* allele (*e^N^* or *e^A^*). F_1_ hybrid females carrying a functional *D. novamexicana ebony* allele (*F*_1_[*e^N^*/*e*^-^]) developed a lighter body color than F_1_ hybrid females carrying a functional *D. americana ebony* allele (F_1_[*e^A^/e*^-^]) (Figure 4B,C vs D,E). These data demonstrate for the first time that functional divergence between the *D. novamexicana* and *D. americana ebony* alleles contributes to divergent body color between these two species.

**Figure 4.**
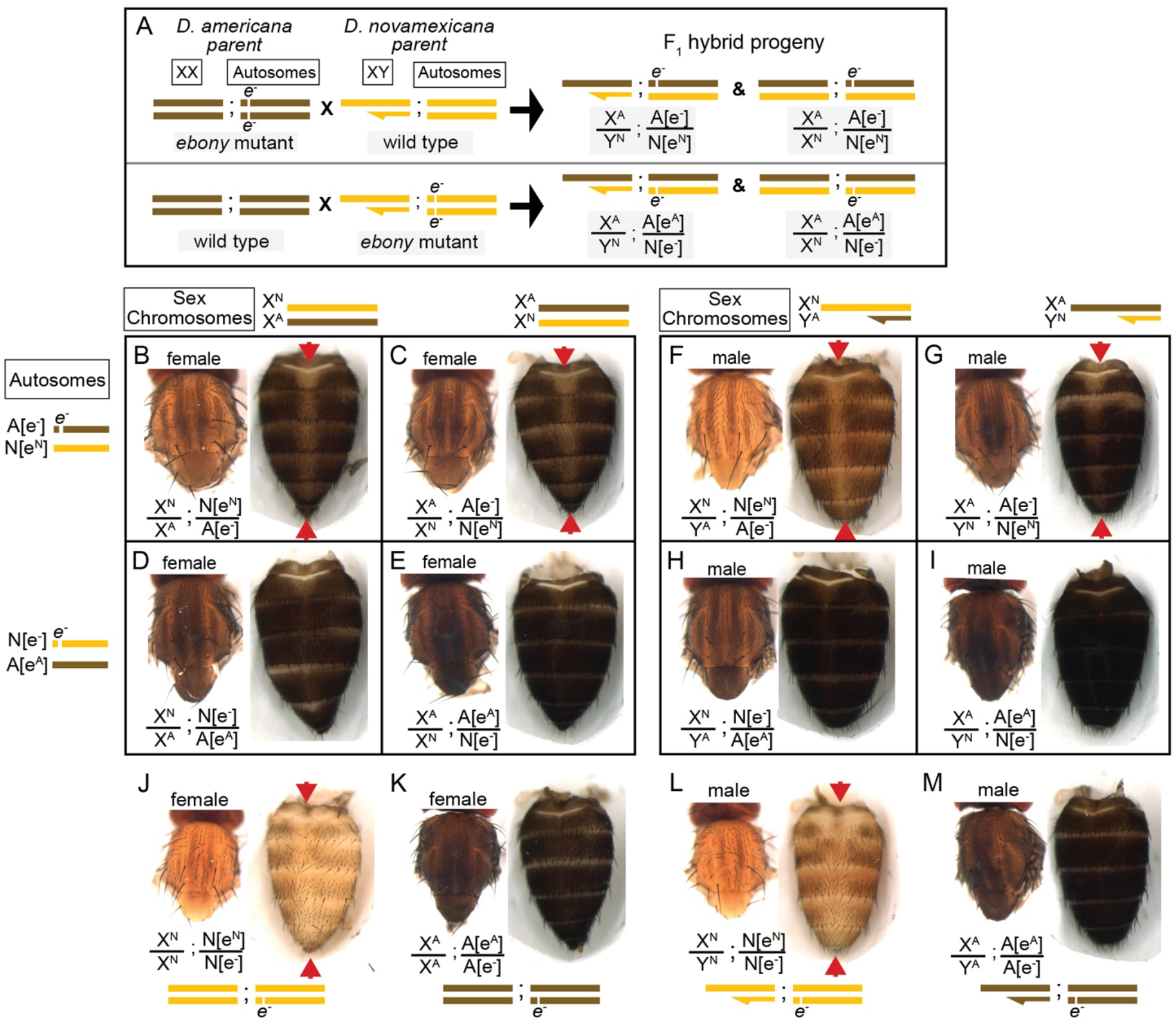
Reciprocal hemizygosity testing shows effects of *ebony* divergence between *D. americana* and *D. novamexicana* on body pigmentation. (**A**) Schematic shows representative sex chromosomes (XX and XY) and autosomes of the parents and progeny of reciprocal hemizygosity crosses, along with the genotypes of the progeny. Although a single autosome is shown for simplicity, these species have five autosomes. Superscript “A” and “N”, as well as brown and yellow colored bars, indicate alleles and chromosomes from *D. americana* and *D. novamexicana*, respectively; *e*^-^ indicates an *ebony* null allele. Although the schematic illustrates the crosses only with *D. americana* as the female parent, the same crosses were performed with sexes of the parental species reversed. (**B-I**) Dorsal thorax and abdomen phenotypes are shown for female (**B-E**) and male (**F-I**) progeny of reciprocal hemizygosity crosses. Genotypes of autosomal and sex chromosomes are shown to the left and above panels **B-I**, respectively, using the same schematic notation as in panel **A**. Individuals in **B**, **C**, **F**, and **G** carry a wild-type copy of *D. novamexicana ebony* allele, whereas individuals in panels **D**, **E**, **H**, and **I** carry a wild-type copy of the *D. americana ebony*. (**J-M**) Dorsal thorax and abdomen phenotypes are shown for female (**J, K**) and male (**L, M**) flies heterozygous for the *ebony* null allele in *D. novamexicana* (**J, L**) and *D. americana* (**K, L**) for comparison to flies shown in panels **B-I**, which also all carry one null and one wild-type *ebony* allele. Red arrowheads in panels **B**, **C**, **F**, **G**, **J**, and **L** highlight the reduced dark pigmentation in the abdomen along the dorsal midline relative to lateral regions.

To determine how *ebony* divergence interacts with divergent loci on the X-chromosome, we also compared the body color of male progeny from these reciprocal crosses. Like the F_1_ hybrid females, these F_1_ hybrid males differ for the parent of origin for their one functional *ebony* allele (*e^A^* or *e^N^*); however, they also differ for the parent of origin of all X-linked genes. Prior work has shown that divergence on the X-chromosome, particularly divergence in non-coding sequences of the *tan* gene, also contributes to differences in body color between *D. novamexicana* and *D. americana* (Wittkopp et al. 2003; Wittkopp et al. 2009). As expected, we found that body color differed between males carrying alternate species’ X chromosomes (Figure 4F vs G and H vs I) as well as between males carrying the same X chromosome but different species’ functional *ebony* alleles (Figure 4F vs H and 3F vs I). Consistent with prior findings demonstrating that divergence in the QTL containing *ebony* explained more of the difference in pigmentation than divergence at X-linked genes, we found that males with functional *D. americana ebony* alleles had the darkest phenotypes, regardless of their X-chromosome genotype (Figure 4F-I).

### 4.3 *ebony* divergence also contributes to a difference in abdominal pigment patterning between *D. novamexicana* and *D. americana*

Although the divergent overall body color is the most striking difference in pigmentation between *D. novamexicana* and *D*. *americana*, there is also a difference in the distribution of pigments along the dorsal midline of the abdomen between these two species (Figure 1). This difference is also visible in individuals of both species heterozygous for an ebony null allele (Figure 4 J-M). Prior work has shown that the absence of dark pigments seen in this region of *D. novamexica* is dominant in F_1_ hybrids to the presence of dark pigments seen in this region of *D. ameriana* (Wittkopp et al. 2003). In addition, genetic mapping of this trait between *D. novamexicana* and *D. virilis* (which has a dark midline region similar to *D. americana)* has shown that the chromosome including *ebony* (chromosome 2) has a large effect on this trait (Spicer 1991). We found that *D. novamexicana ebony* mutants showed even pigmentation across the width of each abdominal segment (Figure 2B), demonstrating that *ebony* is required for the development of lighter pigmentation along the dorsal midline in wild-type D. novamexicana (Figure 2A). In addition, comparing the pigmentation of this abdominal dorsal midline region between F_1_ hybrid flies of both sexes from the reciprocal crosses described above (Figure 4) showed that divergence at *ebony* contributes to this trait difference between *D. novamexicana* and *D. americana*. Specifically, we observed less dark pigments in the dorsal midline region of the abdomen in F_1_ hybrid individuals inheriting the wild-type *D. novamexicana ebony* allele (F_1_[*e^N^*/*e*^-^], Figure 4B,C,F,G) than the *D. americana ebony* allele (F_1_[*e^A^/e*^-^]) (Figure 4D,E,H,I). Males carrying a functional *D. novamexicana ebony* allele (F_1_[*e^N^*/*e*^-^]) showed reduced pigmentation in the dorsal midline relative to the lateral regions regardless of the origin of their X chromosome (Figure 4F,G), indicating that divergent loci on the X-chromosome (including *tan*) do not affect the presence of this phenotype.

### 4.4 Cuticular hydrocarbon profiles differ between *D. americana* and *D. novamexicana* and are affected by *ebony* expression but not *ebony* divergence

*ebony* expression was recently found to affect the relative abundance of cuticular hydrocarbons (CHCs) in *D. melanogaster* (Massey et al. 2019b). In addition, variation in *ebony* expression was also shown to correlate with variation in CHC profiles among natural isolates of *D. melanogaster* (Massey, et al. 2019b). To determine whether *ebony* also effects CHC profiles in *D. novamexicana* and *D. americana*, we extracted CHCs from *D. americana* and *D. novamexicana* female homozygous *ebony* mutant flies as well as females heterozygous for the *ebony* mutant allele. We compared heterozygous individuals (rather than wild-type flies) to mutants homozygous for the *ebony* null allele because they have the same number of functional copies of *ebony* as the reciprocal F_1_ hybrids.

Consistent with a prior report (Bartelt et al. 1986), we found that *D. americana* and *D. novamexicana* produced markedly different CHC profiles. Specifically, we found that CHCs with a chain length of 25 or fewer were only present in *D. novamexicana*, whereas CHCs with a chain length of 31 or greater were only present in *D. americana* (Figure 5A). In both species, the loss of *ebony* function had no qualitative effect on which CHCs were produced by either species, but increased the abundance of some CHCs in both *D. americana* and *D. novamexicana* (Figure 5B,C). *ebony* loss-of-function mutants in *D. melanogaster* were shown to preferentially increase the abundance of long chain CHCs (Massey et al. X), and we observed a similar pattern in *D. americana* (Figure 5D). In *D. novamexicana*, we observed the opposite pattern, however: CHCs with shorter chain lengths showed greater increases in abundance in *ebony* null mutants (Figure 5E). The reason for this difference in how *ebony* affects CHCs in *D. americana* and *D. novamexicana* remains unclear, but might have to do with the different levels of *tan* expression in these two species (Cooley et al. 2012) given that *tan* was also shown to affect CHC profiles in *D. melanogaster* (Massey et al. 2019b).

**Figure 5.**
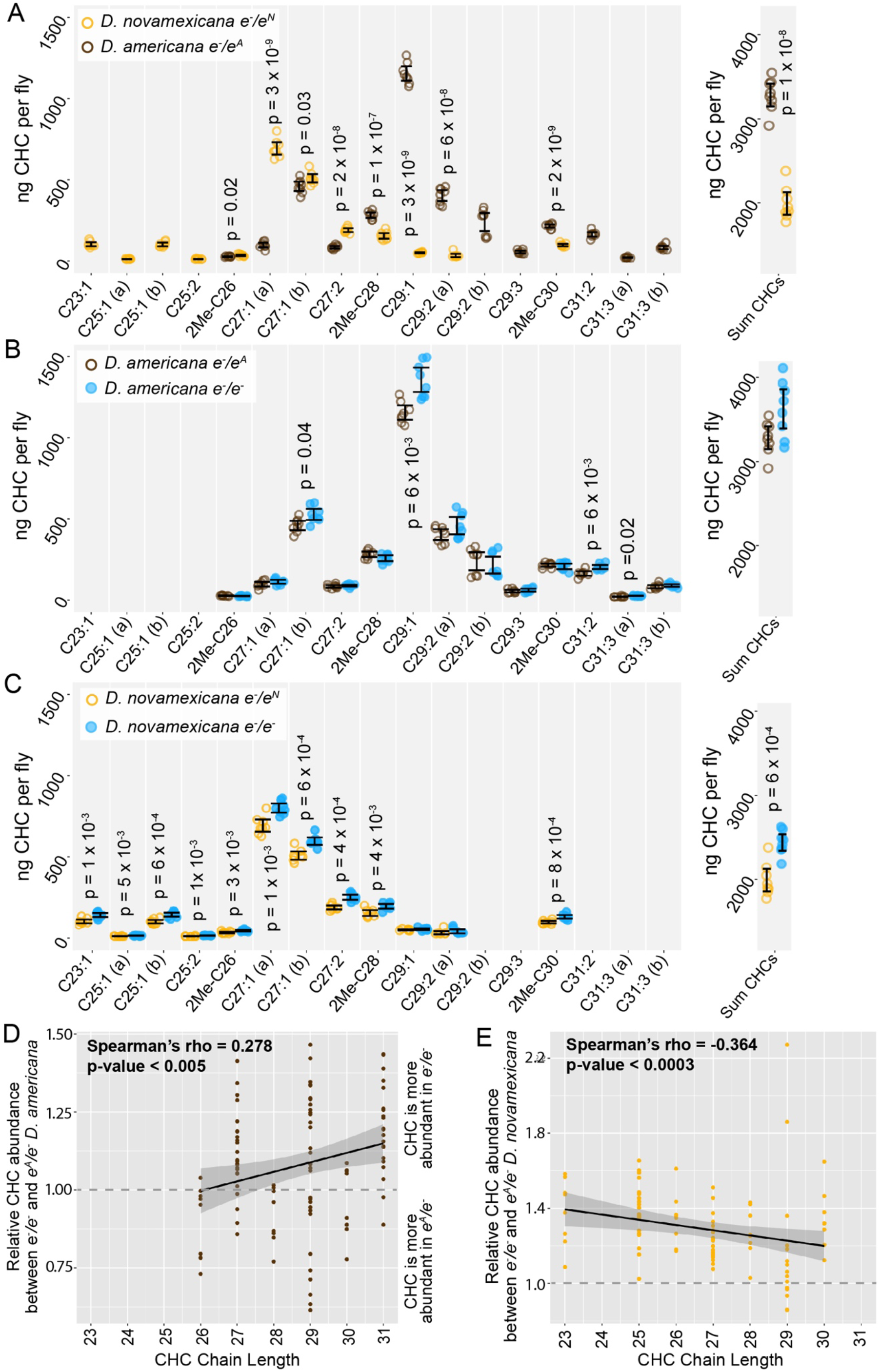
Cuticular hydrocarbons (CHCs) are affected by ebony and differ between D. americana and D. novamexicana. (**A-C**) Abundance of individual CHC compounds (ng/fly) and summed CHCs extracted from female flies are plotted for the following genotypes: (**A**) *D. americana* and *D. novamexicana*, each heterozygous for an *ebony* null (*e*^-^) allele, (**B**) *D. americana* heterozygous and homozygous for an *ebony* null allele, (**C**) *D. novamexicana* heterozygous and homozygous for an *ebony* null allele. Eight biological replicates are shown for each genotype, with error bars representing 95% confidence intervals. For each comparison, the p-value from a Welch’s t-test with a Benjamini-Hochberg multiple test correction *(alpha* = 0.05) is shown when a significant difference in abundance was detected for a CHC present in both genotypes being compared. CHCs are shown from left to right with increasing chain length (represented by “C” followed by the chain length) with double-bond and methyl-branched structures indicated by notations after the colon or before the “C”, respectively. For example, C25:1 represents a 25-carbon monoene, C25:2 represents a 25-carbon diene, and 2Me-C28 represents a 28-carbon alkene with a methyl branch at the second carbon. (**D-E**) Abundance of each CHC in *ebony* null mutants relative to flies heterozygous for the *ebony* null allele is plotted by carbon chain length for (**D**) *D. americana* and (**E**) *D. novamexicana*. Black trendlines in panels **D-E** show linear regressions, with shaded areas representing the standard error and both Spearman’s rho and p-values indicated on each plot.

We also examined the CHC profiles of female F_1_ hybrids produced by crossing *D. americana* females with *D. novamexicana* males. We found that these F_1_ hybrid females showed a CHC profile that was distinct from both species, but more similar to *D. novamexicana* (Figure 6A): it contained some of the short chain CHCs unique to *D. novamexicana* and none of the long chain CHCs unique to *D. americana* (Figure 6A). As seen for both species, eliminating *ebony* function in F_1_ hybrids by making them homozygous for *ebony* null alleles caused an increase in abundance of some CHCs but did not alter which CHCs were present (Figure 6B). Longer chain CHCs were more likely to show increased abundance than shorter chain CHCs (Figure 6C), but this relationship was not as strong as that seen for *D. americana* (Figure 5D). To determine whether divergence between the *D. americana* and *D. novamexicana ebony* alleles affected CHCs profiles, we compared CHCs extracted from females from the reciprocal hemizygosity test. These flies have only one functional *ebony* allele (*D. americana* or *D. novamexicana*) in the F_1_ hybrid genetic background. The CHC profiles from these flies were not significantly different from each other (Figure 6D,E), indicating that allelic divergence at *ebony* does not have a detectable effect on CHCs in this species pair.

**Figure 6.**
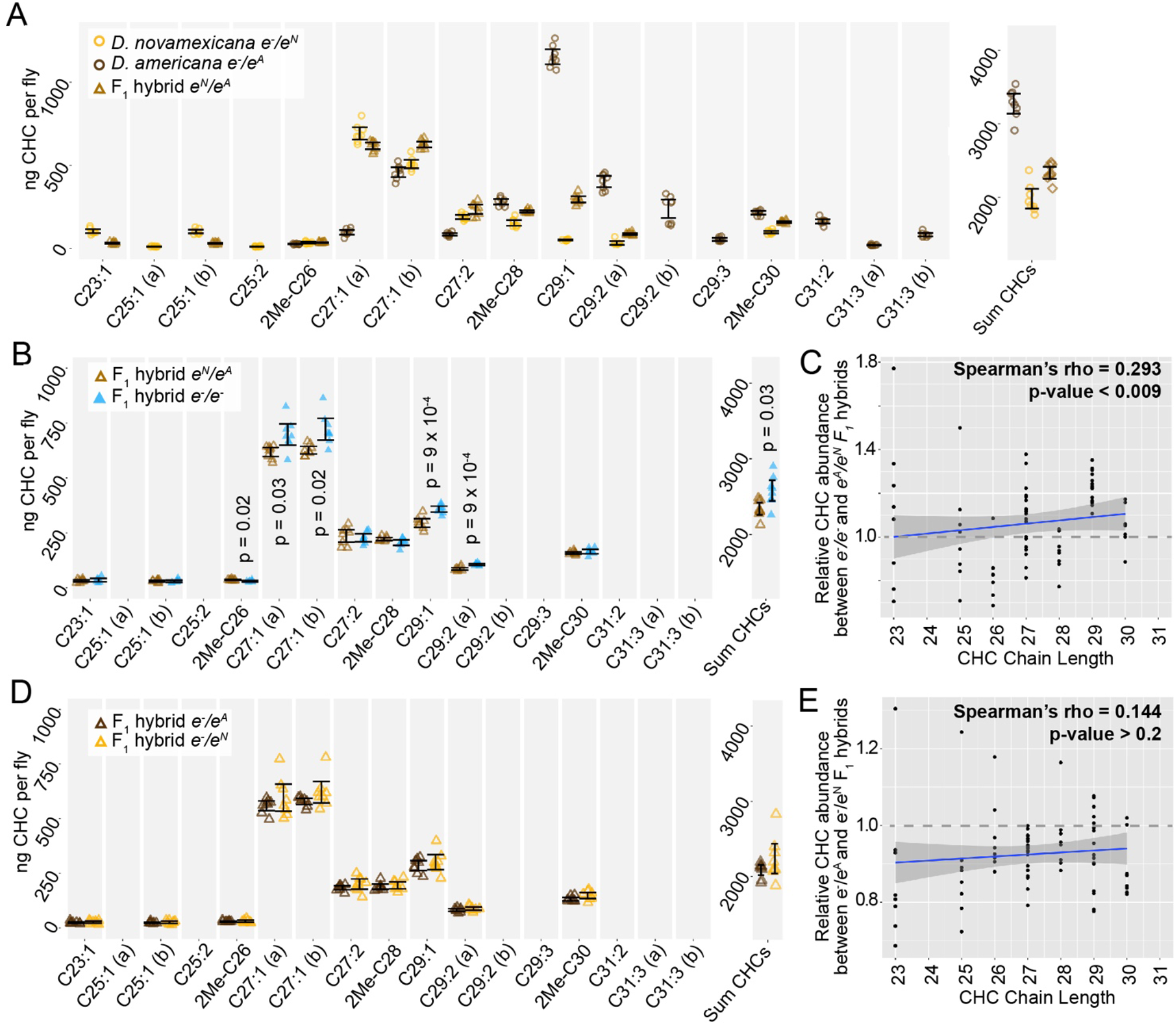
*ebony* does not contribute to divergence of CHCs between *D. americana* and *D. novamexicana*. (**A**) Abundance of individual CHC compounds (ng/fly) and summed CHCs extracted from female flies are plotted for *D. americana* and *D. novamexicana ebony* heterozygotes as well as F_1_ hybrids heterozygous for wild-type alleles of *ebony*. (**B-C**) CHCs from F_1_ hybrids homozygous for *ebony* null alleles are compared to CHCs from F_1_ hybrids with wild-type *D. americana* and *D. novamexicana ebony* alleles, showing the absolute abundance of individual and summed CHC compounds (**B**) as well as the relative abundance of CHCs by carbon chain length (**C**). In panel **B**, p-values are shown from a Welch’s t-test with a Benjamini-Hochberg multiple test correction *(alpha* = 0.05) when a significant difference in abundance was detected for a CHC present in both genotypes. (**D-E**) CHC profiles are plotted for reciprocal F_1_ hybrids that differ only by which wild-type *ebony* allele they carry, either *D. americana* (*e^A^*) or *D. novamexicana* (*e^N^*), with absolute abundance of individual and summed CHCs shown in (**D**) and relative abundance of CHCs by chain length shown in (**E**). No p-values are shown in (**D**) because no CHCs showed a statistically significant difference in abundance between the two F_1_ hybrid genotypes from the reciprocal hemizygosity test (Welch’s t-test with Benjamini-Hochberg multiple test correction, p>0.05 for each CHC). In panels **C** and **E**, blue trendlines show linear regressions, with shaded areas representing the standard error and both Spearman’s rho and p-values indicated on each plot. In all panels, data from eight replicate flies is shown for each genotype.

## 5 Conclusions

Identifying the genes responsible for phenotypic differences between species remains a significant challenge for evolutionary biology. This task is especially challenging when a gene contributing to phenotypic divergence is located in a region of the genome inverted between species, which precludes recombination-based mapping. Such is the case for the *ebony* gene in *D. americana* and *D. novamexicana*. Prior work suggested that *ebony* might contribute to differences in overall body color between these two species (Wittkopp et al. 2009; Cooley et al. 2012), but its location in an inversion made it difficult to directly test this hypothesis. In this study, we overcame this hurdle by using CRISPR/Cas9 genome editing to generate null mutants for *ebony* in *D. americana* and *D. novamexicana*, and then using these mutants to perform a reciprocal hemizygosity test (Stern 2014), which directly compares the effects of the two species’ alleles on pigmentation. We found that divergence at *ebony* does indeed contribute to differences in body color between *D. americana* and *D. novamexicana*.

Characterizing the phenotypes of *D. americana* and *D. novamexicana ebony* mutants, as well as flies from the reciprocal hemizygosity test, also identified effects of *ebony* on other phenotypes. For example, we found that differences in the activity of *ebony* alleles between *D. americana* and *D. novamexicana* are responsible for the absence of dark pigmentation seen along the dorsal abdominal midline of *D. novamexicana* but not *D. americana*. This trait has previously been described as derived in *D. novamexicana* (Spicer 1991); however, we see a similar dorsal midline lightening in at least some lines of *D. lummei* (see Figure 1), another member of the virilis group, suggesting that the dorsal midline activity of *ebony* existed prior to the divergence of *D. americana* and *D. novamexicana*. An unexpected change in pupal pigment patterning was also seen in *D. american*a and *D. novamexicana ebony* null mutants. Although *ebony* is known to affect pupal case development in *D. melanogaster* (Sherald 1980), its loss causes a pale white pupa color rather than the dark pigmentation we see in *D. americana* and *D. novamexicana ebony* null mutants. Because *ebony* is required for the production of yellow pigments, the dark markings seen in *ebony* mutant pupal cases likely result from expression of an enzyme required for synthesis of dark pigments, such as *tan*. Finally, we found that *ebony* null mutants showed significant changes in the abundance of some CHCs in each species, but divergence of *ebony* did not contribute to differences in the CHC profiles seen between species. These observations illustrate how *cis*-regulatory changes can cause divergence of some, but not all, traits affected by a pleiotropic gene.

Observations reported in this work were made possible by the ability to manipulate the *D. americana* and *D. novamexicana* genomes with CRISPR/Cas9 genome editing. While this technology has great potential for allowing functional hypothesis testing in species that have not historically been considered genetic model systems, this work was not always straightforward. We hope that the detailed descriptions of our genome editing efforts provided in the Materials and Methods section of this paper will be helpful for other researchers striving to manipulate the genomes of non-model species.

## 6 Author Contributions

AML and PJW conceived of the experiments. AML, PS, and ZW performed the experiments. HC and PWJ provided funding, advice, and oversight. AML and PJW wrote the manuscript.

## 7 Funding

This work was funded by the National Institutes of Health (Grant No. 1R35GM118073 and Grant No. 1R01GM089736) awarded to PJW; National Science Foundation Graduate Research Fellowship Program (Grant No: DGC 1256260) and National Institute of Health training grant: “Michigan Predoctoral Training in Genetics” (Grant No: T32GM00754) to AML; and startup funding provided by Michigan State University AgbioResearch to HC.

## 8 Conflict of Interest Statement

The authors declare that the research was conducted in the absence of any commercial or financial relationships that could be construed as a potential conflict of interest.

## 9 Acknowledgements

We thank Arnaud Martin (George Washington University) as well as Kathy Vaccarro and other members of Sean Carroll’s laboratory (University of Wisconsin) for advice on CRISPR/Cas9 genome editing and *Drosophila* injections, respectively; Hannah McConnell, Aida de la Cruz, and Harmit Malik (Fred Hutchinson Cancer Research Center) for sharing their experience working with the *nanos* promoter in *Drosophila virilis;* and the Bloomington Drosophila Stock Center as well as the National Drosophila Species Stock Center for maintaining and providing fly stocks

## References

Ahmed-Braimah, Y.H. and McAllister, B.F. (2012). Rapid evolution of assortative fertilization between recently allopatric species of *Drosophila*. Int. J. Evol. Biol. 2012, 285468. doi: 10.1155/2012/285468.

Ahmed-Braimah, Y.H. and Sweigart, A.L. (2015). A single gene causes an interspecific difference in pigmentation in *Drosophila*. Genetics 200, 331–42. doi: 10.1534/genetics.115.174920.

Bartelt, R.J., Armold, M.T., Schaner, A.M., and Jackson, L.L. (1986). Comparative analysis of cuticular hydrocarbons in the *Drosophila virilis* species group. Comp. Biochem. Physiol. -- Part B Biochem. 83, 731–42. doi: 10.1016/0305-0491(86)90138-0.

Bassett, A.R., Tibbit, C., Ponting, C.P., and Liu, J.-L. (2013). Highly Efficient Targeted Mutagenesis of *Drosophila* with the CRISPR/Cas9 System. Cell Rep. 4, 220–28. doi: 10.1016/j.celrep.2013.06.020.

Benjamini, Y. and Hochberg, Y. (1995). Controlling the false discovery rate: a practical and powerful approach to multiple testing. J. R. Stat. Soc. Ser. B 57, 289–300. https://www.jstor.org/stable/2346101.

Caletka, B.C. and McAllister, B.F. (2004). A genealogical view of chromosomal evolution and species delimitation in the *Drosophila virilis* species subgroup. Mol. Phylogenet. Evol. 33, 664–70. doi: 10.1016/j.ympev.2004.08.007.

Carroll, S.B. (2005). Evolution at two levels: on genes and form. PLoS Biol. 3, e245. doi: 10.1371/journal.pbio.0030245.

Chung, H. and Carroll, S.B. (2015). Wax, sex and the origin of species: Dual roles of insect cuticular hydrocarbons in adaptation and mating. BioEssays, 822–30. doi: 10.1002/bies.201500014.

Chung, H., Loehlin, D.W., Dufour, H.D., Vaccarro, K., Millar, J.G., and Carroll, S.B. (2014). A single gene affects both ecological divergence and mate choice in *Drosophila*. Science 343, 1148–51. doi: 10.1126/science.1249998.

Cooley, A.M., Shefner, L., McLaughlin, W.N., Stewart, E.E., and Wittkopp, P.J. (2012). The ontogeny of color: developmental origins of divergent pigmentation in *Drosophila americana* and *D. novamexicana*. Evol. Dev. 14, 317–25. doi: 10.1111/j.1525-142X.2012.00550.x.

Holtzman, S., Miller, D., Eisman, R., Kuwayama, H., Niimi, T., and Kaufman, T. (2010). Transgenic tools for members of the genus *Drosophila* with sequenced genomes. Fly (Austin). 4, 349–62. doi: 10.4161/fly.4.4.13304.

Horn, C. and Wimmer, E.A. (2000). A versatile vector set for animal transgenesis. Dev. Genes Evol. 210, 630–37. doi: 10.1007/s004270000110.

Koch, P.B., Behnecke, B., Weigmann-Lenz, M., and Ffrench-Constant, R.H. (2000). Insect pigmentation: Activities of β-alanyldopamine synthase in wing color patterns of wild-type and melanic mutant swallowtail butterfly *Papilio glaucus*. Pigment Cell Res. 13, 54–58. doi: 10.1111/j.0893-5785.2000.130811.x.

Kronforst, M.R., Barsh, G.S., Kopp, A., Mallet, J., Monteiro, A., Mullen, S.P., Protas, M., Rosenblum, E.B., Schneider, C.J., and Hoekstra, H.E. (2012). Unraveling the thread of nature’s tapestry: the genetics of diversity and convergence in animal pigmentation. Pigment Cell Melanoma Res. 25, 411–33. doi: 10.1111/j.1755-148X.2012.01014.x.

Liu, M., Rehman, S., Tang, X., Gu, K., Fan, Q., Chen, D., and Ma, W. (2018). Methodologies for Improving HDR Efficiency. Front. Genet. 9, 691. doi: 10.3389/fgene.2018.00691.

Massey, J. H. and Wittkopp, P.J. (2016). The genetic basis of pigmentation differences within and between *Drosophila* species. In Curr. Top. Dev. Biol., 119:27–61. doi: 10.1016/bs.ctdb.2016.03.004.

Massey, Jonathan H., Chung, D., Siwanowicz, I., Stern, D.L., and Wittkopp, P.J. (2019a). The *yellow* gene influences *Drosophila* male mating success through sex comb melanization. eLife 8, 1–20. doi: 10.7554/eLife.49388.

Massey, J. H., Akiyama, N., Bien, T., Dreisewerd, K., Wittkopp, P.J., Yew, J.Y., and Takahashi, A. (2019b). Pleiotropic effects of *ebony* and *tan* on pigmentation and cuticular hydrocarbon composition in *Drosophila melanogaster*. Front. Physiol. 10, 518. doi: 10.3389/fphys.2019.00518.

Miller, D.F.B., Holtzman, S.L., and Kaufman, T.C. (2002). Customized microinjection glass capillary needles for P-element transformations in *Drosophila melanogaster*. Biotechniques 33, 366–75. doi: 10.2144/02332rr03.

Morales-Hojas, R., Vieira, C.P., and Vieira, J. (2008). Inferring the evolutionary history of *Drosophila americana* and *Drosophila novamexicana* using a multilocus approach and the influence of chromosomal rearrangements in single gene analyses. Mol. Ecol. 17, 2910–26. doi: 10.1111/j.1365-294X.2008.03796.x.

Nappi, A.J. and Christensen, B.M. (2005). Melanogenesis and associated cytotoxic reactions: Applications to insect innate immunity. Insect Biochem. Mol. Biol. 35, 443–59. doi: 10.1016/j.ibmb.2005.01.014.

Patterson, J.T. and Stone, W.S. (1949). The relationship of *novamexicana* to the other members of the *virilis* group. In Univ. Texas Publ., 4920:7–17.

Port, F., Chen, H.M., Lee, T., and Bullock, S.L. (2014). Optimized CRISPR/Cas tools for efficient germline and somatic genome engineering in *Drosophila*. Proc. Natl. Acad. Sci. U. S. A. 111, E2967–76. doi: 10.1073/pnas.1405500111.

Rebeiz, M. and Williams, T.M. (2017). Using *Drosophila* pigmentation traits to study the mechanisms of cis-regulatory evolution. Curr. Opin. Insect Sci. 19, 1–7. doi: 10.1016/j.cois.2016.10.002.

Sherald, A.F. (1980). Sclerotization and coloration of the insect cuticle. Experientia 36, 143–46. doi: 10.1007/BF01953696.

Spicer, G.S. (1991). The genetic basis of a species-specific character in the Drosophila virilis species group. Genetics 128, 331–37. http://www.ncbi.nlm.nih.gov/pubmed/2071018.

Stern, D.L. (2014). Identification of loci that cause phenotypic variation in diverse species with the reciprocal hemizygosity test. Trends Genet. 30, 547–54. doi: 10.1016/j.tig.2014.09.006.

Stern, D.L., Crocker, J., Ding, Y., Frankel, N., Kappes, G., Kim, E., Kuzmickas, R., Lemire, A., Mast, J.D., and Picard, S. (2017). Genetic and transgenic reagents for *Drosophila simulans, D. mauritiana, D. yakuba, D. santomea*, and *D. virilis*. G3 (Bethesda). 7, 1339–47. doi: 10.1534/g3.116.038885.

Suh, J. and Jackson, F.R. (2007). *Drosophila ebony* activity is required in glia for the circadian regulation of locomotor activity. Neuron 55, 435–47. doi: 10.1016/j.neuron.2007.06.038.

Takahashi, A. (2013). Pigmentation and behavior: potential association through pleiotropic genes in *Drosophila*. Genes Genet. Syst. 88, 165–74. http://www.ncbi.nlm.nih.gov/pubmed/24025245.

Thurmond, J., Goodman, J.L., Strelets, V.B., Attrill, H., Gramates, L.S., Marygold, S.J., Matthews, B.B., Millburn, G., Antonazzo, G., Trovisco, V., Kaufman, T.C., Calvi, B.R., and FlyBase Consortium. (2019). FlyBase 2.0: the next generation. Nucleic Acids Res. 47, D759–65. doi: 10.1093/nar/gky1003.

True, J.R. (2003). Insect melanism: the molecules matter. Trends Ecol. Evol. 18, 640–47. doi: 10.1016/j.tree.2003.09.006.

True, J.R., Yeh, S.D., Hovemann, B.T., Kemme, T., Meinertzhagen, I. a, Edwards, T.N., Liou, S.R., Han, Q., and Li, J. (2005). *Drosophila tan* encodes a novel hydrolase required in pigmentation and vision. PLoS Genet. 1, e63. doi: 10.1371/journal.pgen.0010063.

Wittkopp, P. J., Stewart, E.E., Arnold, L.L., Neidert, A.H., Haerum, B.K., Thompson, E.M., Akhras, S., Smith-Winberry, G., and Shefner, L. (2009). Intraspecific polymorphism to interspecific divergence: genetics of pigmentation in *Drosophila*. Science. 326, 540–44. doi: 10.1126/science.1176980.

Wittkopp, P J, Smith-Winberry, G., Arnold, L.L., Thompson, E.M., Cooley, a M., Yuan, D.C., Song, Q., and McAllister, B.F. (2011). Local adaptation for body color in *Drosophila americana*. Heredity (Edinb). 106, 592–602. doi: 10.1038/hdy.2010.90.

Wittkopp, Patricia J, and Beldade, P. (2009). Development and evolution of insect pigmentation: genetic mechanisms and the potential consequences of pleiotropy. Semin. Cell Dev. Biol. 20, 65–71. doi: 10.1016/j.semcdb.2008.10.002.

Wittkopp, Patricia J, Carroll, S.B., and Kopp, A. (2003). Evolution in black and white: genetic control of pigment patterns in *Drosophila*. Trends Genet. 19, 495–504. doi: 10.1016/S0168-9525(03)00194-X.

Wittkopp, Patricia J, True, J.R., and Carroll, S.B. (2002). Reciprocal functions of the Drosophila yellow and ebony proteins in the development and evolution of pigment patterns. Development 129, 1849–58. http://www.ncbi.nlm.nih.gov/pubmed/11934851.

Wittkopp, Patricia J, Williams, B.L., Selegue, J.E., and Carroll, S.B. (2003). *Drosophila* pigmentation evolution: divergent genotypes underlying convergent phenotypes. Proc. Natl. Acad. Sci. U. S. A. 100, 1808–13. doi: 10.1073/pnas.0336368100.

Wray, G.A., Hahn, M.W., Abouheif, E., Balhoff, J.P., Pizer, M., Rockman, M. V, and Romano, L.A. (2003). The evolution of transcriptional regulation in eukaryotes. Mol. Biol. Evol. 20, 1377–1419. doi: 10.1093/molbev/msg140.

