## Supplemental_figs_tables for "*ebony* affects pigmentation divergence and cuticular hydrocarbons in *Drosophila americana* and *D. novamexicana*"

1 **Supplemental Materials**

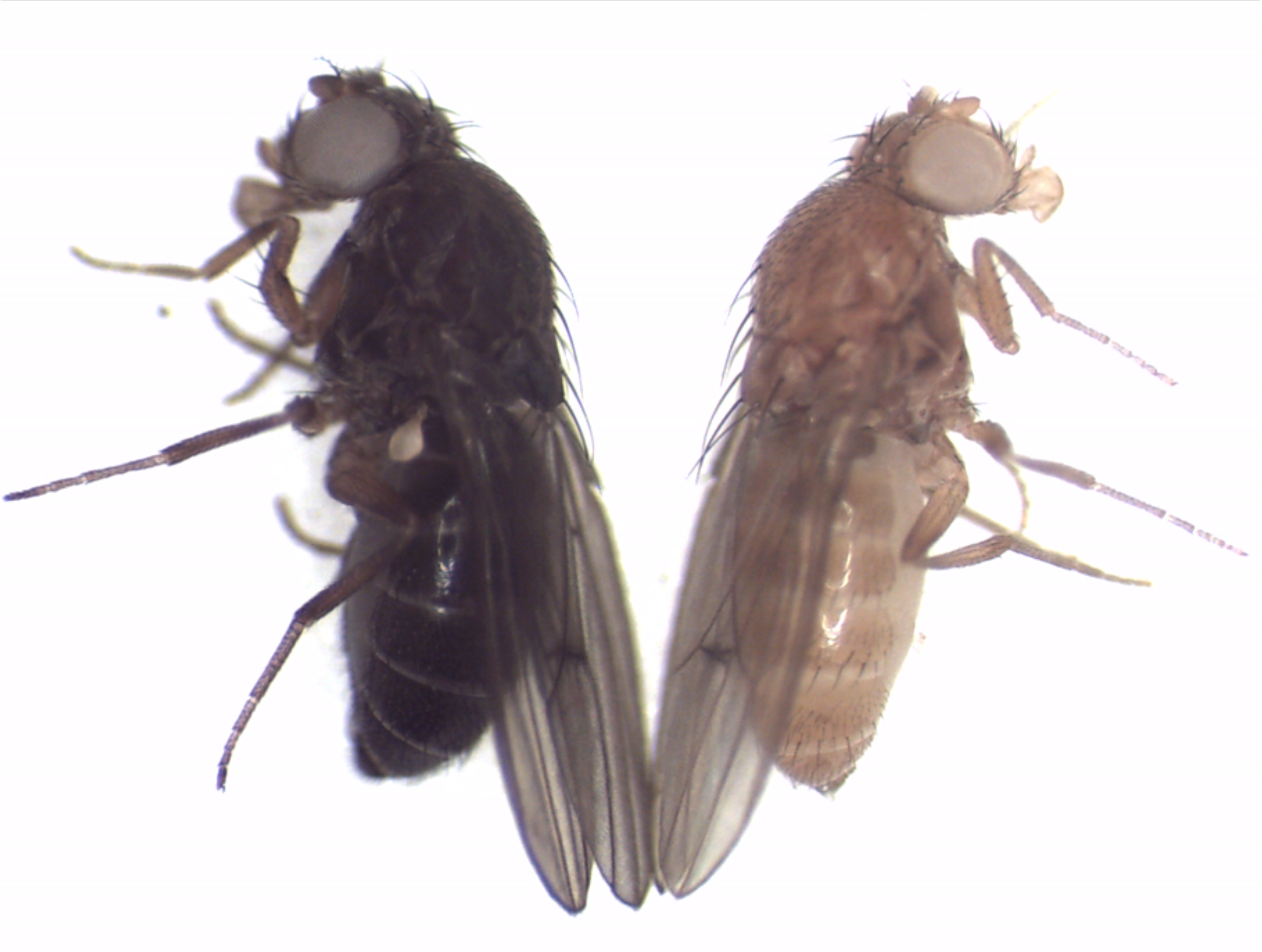

**Supplementary Figure 1. Mutations in the *D. novamexicana* and *D. americana white* genes cause white-eyed phenotype.** Photograph shows *D. americana white* mutant adult male (left) alongside *D. novamexicana white* mutant mutant adult male (right).

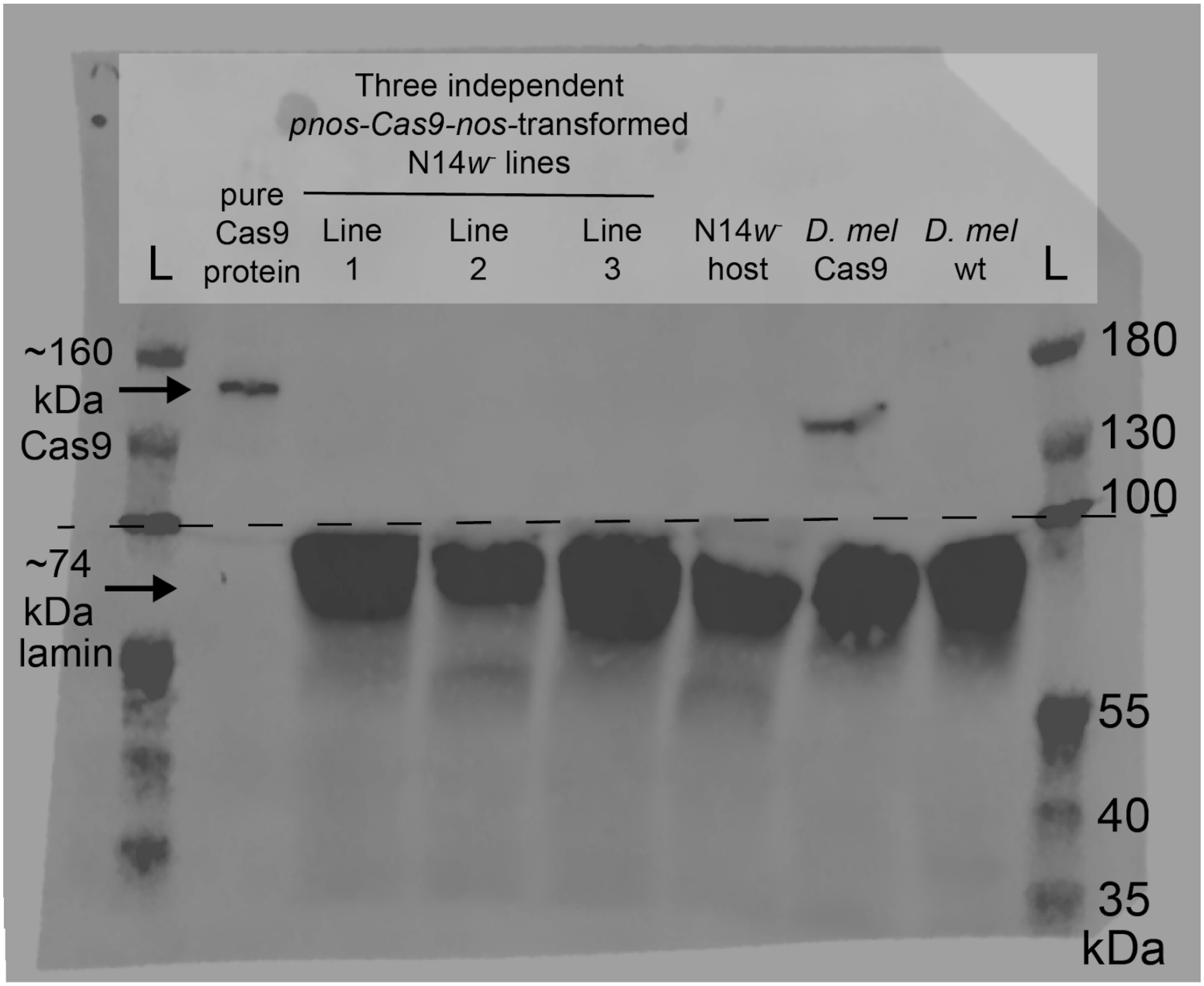


**Supplementary Figure 2. *nanos* promoter failed to drive expression of Cas9 in transgenic *D. novamexicana*.** Western blotting showed that three independent insertions of a transgene expected to express Cas9 protein in the germline under the control of the *nanos* promoter failed to produce detectable levels of Cas9. From left to right, lanes contained a sample of pure Cas9 protein obtained commercially (a positive control) and then total protein extracts from ovaries of sexually mature adult female flies from three lines of *D. novamexicana* with independent insertions of the piggyBac transgene (lines 1-3), the *D. novamexicana* N14 *white* mutant (*w*$^{-}$) host line transgenes were injected into (a negative control), a line of *D. melanogaster* expressing Cas9 in the germline (a second positive control), and a wild-type (wt) of line of *D. melanogaster* (a second negative control). “L” represents the ladder, or molecular weight marker (PageRuler prestained protein ladder). The dotted black line shows where the membrane was cut prior to incubation with primary antibodies during the western blotting procedure; the top half was incubated with anti-Cas9 antibodies whereas the bottom half was incubated with anti-Lamin antibodies. The two halves were realigned by hand for imaging, using the shape of the cut and the ladder staining as a guide. Relative intensity of the protein detected with the antibody against Lamin estimate the relative amounts of total protein loaded per lane. An un-annotated image of this blot is shown in Supplementary Figure 4.

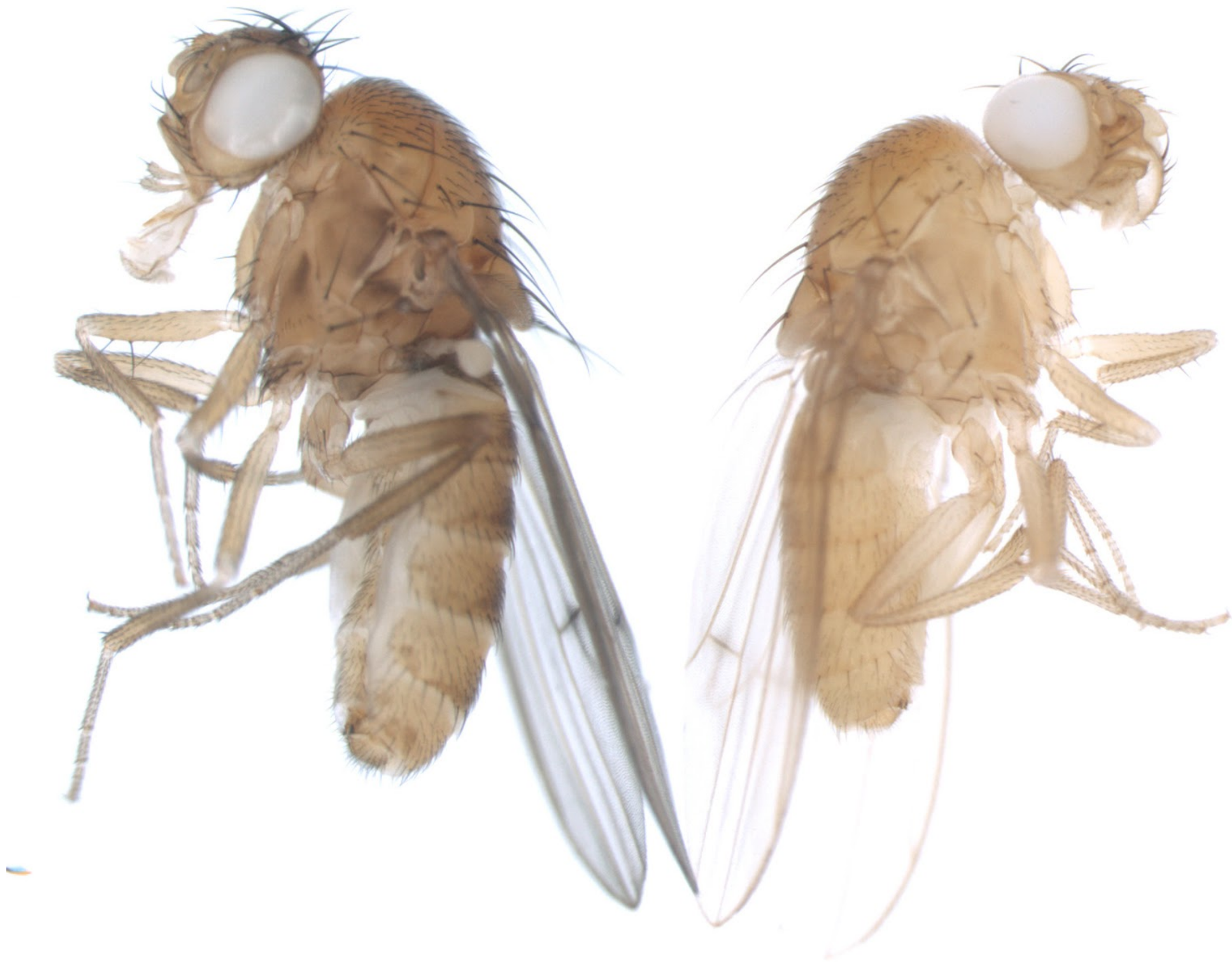

**Supplementary Figure 3. Mutation in the *D. novamexicana yellow* gene causes visible changes in body pigmentation.** Photograph shows *D. novamexicana white* mutant adult male (left) alongside *D. novamexicana white, yellow* mutant adult male (right). Consistent with *yellow* null phenotypes in other species, *D. novamexicana* individuals identified as *yellow* mutants displayed a complete lack of black pigmentation on body, wings, and bristles. Phenotypically *yellow* males were observed only in crosses where the female N14 *nos-Cas9-nos* transgenic parent was injected with Cas9 protein along with sgRNAs targeting conserved sites in *yellow* exon 1.

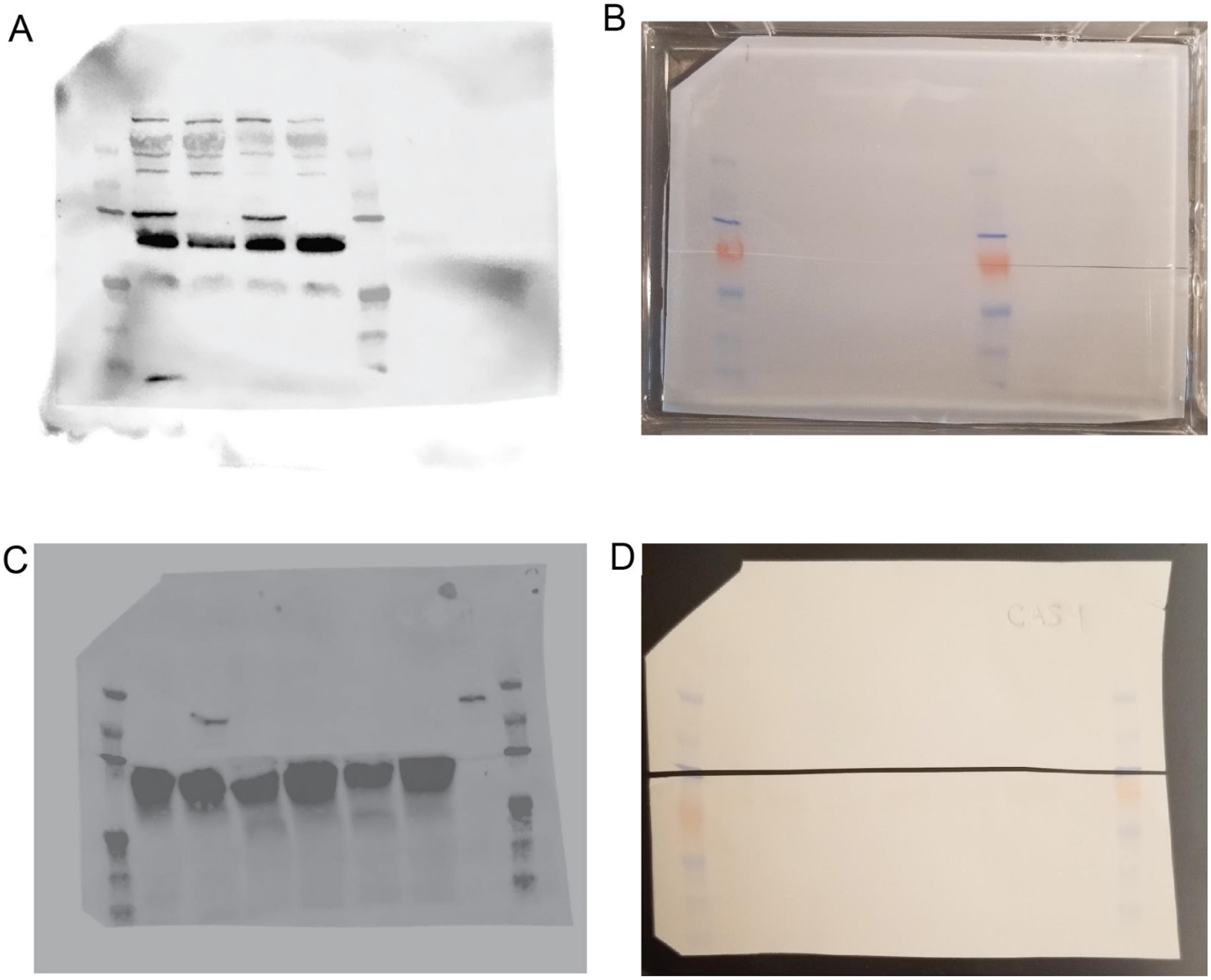


**Supplementary Figure 4. Uncropped and un-annotated western blot images and membranes.** (**A-B**) Images show combined enhanced chemiluminescence (ECL) and 700nm ladder channels (**A**) and photograph of the blocked and cut membrane (**B**) from Ebony western blot in Figure 3B. (**C-D**) Images show combined enhanced chemiluminescence (ECL) and 700nm ladder channels (**C**) and photograph of the cut membrane (**D**) from Cas9 western blot in Supplementary Figure 2.

**Supplementary Table 1. Oligonucleotides used to generate sgRNAs for CRISPR/Cas9 genome editing and *nos-Cas9-nos* transgene.**

| Name | Purpose | Sequence |
|---|---|---|
| IVT_Common_Rev | Common reverse primer for generating all DNA templates for in-vitro transcription of sgRNAs | AAAAGCACCGACTCGGTGCCACTTTTTCAAGTTGATAACGGACTAGCCTTATTTTAACTTGCTATTTCTAGCTCTAAAAC |
| IVT_e_ex2A_Fw | Forward primer for generating DNA template for in-vitro transcription of sgRNA targeting one site in *ebony* exon 2. Target site in bold. | GAAATTAATACGACTCACTATAG**GCTGCGCCATGCCGACAAGG**GTTTTAGAGCTAGAAATAGC |
| IVT_e_ex2B_Fw | | GAAATTAATACGACTCACTATAG**GCTCATTTATCAGCAGGAAC**GTTTTAGAGCTAGAAATAGC |
| IVT_e_ex2C_Fw | | GAAATTAATACGACTCACTATAGG**TGCCGCCCGTCTCCTTGTGG**GTTTTAGAGCTAGAAATAGC |
| IVT_e_ex2D_Fw | | GAAATTAATACGACTCACTATAGG**CAGCGATGGGGACTTCATCG**GTTTTAGAGCTAGAAATAGC |
| IVT_e_ex2E_Fw | | GAAATTAATACGACTCACTATAGG**GTTTGCATGCAACCGTCGG**GTTTTAGAGCTAGAAATAGC |
| IVT_w_ex2_Fw | Forward primer for generating DNA template for in-vitro transcription of | GAAATTAATACGACTCACTATAG**GTCCACATTGTGCCACGCAT**GTTTTAGAGCTAGAAATAGC |

| | | |
|---|---|---|
| IVT_w_ex3_Fw | sgRNA targeting one site in either *white* exon 2 or 3. Target site in bold. | GAAATTAATACGACTCACTATAG**GACCACGCTGCTGAATGCCC**GTTTTAGAGCTAGAAATAGC |
| IVT_yA_Fw | Forward primer for generating DNA template for in-vitro transcription of sgRNA targeting one site in *yellow* exon 1. Target site in bold. | GAAATTAATACGACTCACTATAGG**AAGGAGCAGGCGATCGCCAG**GTTTTAGAGCTAGAAATAGC |
| IVT_yB_Fw | | GAAATTAATACGACTCACTATAGG**CCGCAGAACGGCCTGCCCGT**GTTTTAGAGCTAGAAATAGC |
| IVT_yC_Fw | | GAAATTAATACGACTCACTATAGG**TCTTTGTCACAGTGCCGCGC**GTTTTAGAGCTAGAAATAGC |
| nosCas9_GA_Fw | Amplify *nos-Cas9-nos* transgene out of p*nos-Cas9-nos* (Addgene #62208) with tailed primers to assemble into pBac{3XP3-ECFPafm}. | TTCGAATGGCCATGGGACGTCGACCGGATTTCACTGGAACTAGGCTAG |
| nosCas9_GA_Rev | | ATATAGGGCCCGGGTTATAATTACCCGAGACCGTGACCTACATCG |

**Supplementary File 1. Raw data measuring CHC abundance**

**Supplementary File 2. R code used for analyzing CHC data**
